# The *Drosophila* gene *smoke alarm* regulates nociceptor-epidermis interactions and thermal nociception behavior

**DOI:** 10.1101/2021.05.12.443649

**Authors:** Stephanie E. Mauthner, Katherine H. Fisher, W. Daniel Tracey

## Abstract

The detection and processing of noxious sensory input depends on the proper growth and function of nociceptor sensory neurons in the peripheral nervous system. In *Drosophila melanogaster*, the class IV (cIV) multidendritic dendritic arborization (md-da) neurons detect noxious stimuli through their highly branched dendrites that innervate the epidermis of the larval body wall. Here, we describe requirements of a previously uncharacterized gene named *smoke alarm* (*smal*), a discoidin domain receptor, in cIV md-da dendrite morphogenesis and nociception behavior. We find that *smal* mutant larvae exhibit thermal hyperalgesia that is fully rescued with a BAC transgene containing *smal*. Consistent with this phenotype, a *smal* reporter gene was expressed in nociceptors and other peripheral sensory neurons. Smal::GFP protein localized to punctate structures throughout the cIV md-da neurons. We further show that *smal* loss-of-function results in reduced nociceptor dendrite branching. Interestingly, mammalian homologues of *smal* act as collagen receptors, and we find that *smal* mutant dendrites showed an increase in epidermal cell ensheathment relative to animals that are wild type for *smal*. Based on this phenotype we propose that Smal protein function is required for attachment of dendrites to the extracellular matrix (ECM) and the loss of this activity results in thermal hyperalgesia.

## Introduction

Nociception refers to sensing of noxious stimuli, which have the potential to cause tissue damage. This is accomplished using nociceptors—sensory neurons that are specialized to sense noxious stimuli in the environment. Nociceptors may be polymodal and respond to a variety of noxious stimuli (i.e. thermal, mechanical, or chemical signals) or they may be specialized to detect a specific insult. The sensory transduction machinery in nociceptors is specifically activated by noxious stimuli in the environment and the signals are propagated to higher order neurons in the central nervous system that produce defensive behavioral outputs. While molecular mechanisms of nociception and pain have largely been studied in mammals, progress has also been made from studies of invertebrates like fruit files (Tracey et al., 2003; Hwang et al., 2007; Neely et al., 2010; Zhong et al., 2010; Milinkeviciute et al., 2012; Zhong et al., 2012; Honjo et al., 2016; Jo et al., 2017; Khuong et al., 2019).

The larvae of *Drosophila melanogaster* have been developed as a powerful model to study pain-signaling mechanisms. *Drosophila* larvae have stereotyped behavioral responses to painful stimuli. When presented with a noxious stimulus, larvae exhibit a highly stereotyped behavioral response, termed nocifensive escape locomotion, or rolling in which larvae rotate around the long body axis in a corkscrew-like motion (Tracey et al., 2003). Genetics screens for mutants and genes required for this behavior makes *Drosophila* larvae a robust system for identifying the molecular mechanisms of nociception. Moreover, many fly nociception genes are conserved in mammals (Woolf et al., 1997; Kang et al., 2010; Neely et al., 2010; Honjo et al., 2014; Jo et al., 2017),

In this system, multidendritic dendritic arborization (md-da) peripheral sensory neurons (Grueber *et al*., 2002) have been identified as the larval nociceptors (Hwang et al., 2007; Hu et al., 2017). The class IV (cIV) md-da neurons are the major polymodal larval pain-sensing neurons (Hwang et al., 2007). These highly branched neurons extensively cover the larval body wall, and are both necessary and sufficient for triggering rolling behavior (Hwang et al., 2007). In addition to cIV neurons other md-da neurons (Class II and Class III) also participate in mechanical nociception responses (Hu et al., 2017).

Laser capture microdissection and microarray analysis has identified genes enriched in larval nociceptors (Mauthner et al., 2014). The nociceptor-enriched genes were further studied with a cIV targeted thermal nociception RNAi screen to identify genes that are functionally required for thermal nociception behaviors. Some of the identified genes were found to display dendritic morphology changes (Honjo *et al*., 2016). This screen uncovered *CG34380*, a previously uncharacterized gene that was named *smoke alarm* (*smal*), as RNAi targeting the gene caused increased sensitivity to noxious heat (Honjo et al., 2016). Additionally, *smal* knockdown resulted in a significant increase in cIV dendrite coverage along the larval body wall (Honjo *et al*., 2016).

*smoke alarm* encodes a discoidin domain receptor (DDR) protein. Mammalian homologues of Smal are type-I transmembrane receptor tyrosine kinases (RTKs) that are activated by binding to collagen. To date, previous research on DDRs has been focused on its role in cancer in humans, and axon guidance and regeneration in *C. elegans* (Unsoeld et al., 2013; Leitinger, 2014; Hisamoto et al., 2016). While transcript expression of the *Ddr* gene family has been previously identified in (Chiu et al., 2014) and *Drosophila* nociceptive neurons (Honjo et al., 2016), the function of DDR proteins in nociception are unknown.

*Drosophila* md-da neuron dendrites are sandwiched between the epidermis and its basal lamina extracellular matrix (ECM). Interactions between these neurons, the epidermis, and the ECM are crucial for proper neuronal growth and sensory function (Han et al., 2012; Kim et al., 2012; Tenenbaum et al., 2017; Jiang et al., 2019(Yasunaga et al., 2010)). The nociceptor-ECM interactions are critical to the avoidance of redundant innervation via proper dendritic patterning and growth (Han et al., 2012; Kim et al., 2012; Meltzer et al., 2016; Tenenbaum et al., 2017; Hoyer et al., 2018; Jiang et al., 2019). Several studies have demonstrated that nociceptors are physically attached to the basal lamina through integrin proteins, which bind to laminin proteins in the ECM (Han et al., 2012; Kim et al., 2012). Mutations in the alpha and beta integrin subunits, *multiple edematous wings (mew)* and *myospheroid (mys)*, cause cIV dendrites to detach from the basal lamina and as a result, cIV dendrites show increased ensheathment by the epidermis. Some ensheathment is important for stabilization of dendritic branches (Tenenbaum et al., 2017). However, abnormally high levels of ensheathment result in failures of dendrite to self-avoid, leading to dendritic crossovers and aberrant dendrite branching (Han et al., 2012; Kim et al., 2012). Additionally, blocking dendrite ensheathment via overexpression of integrin proteins or knockdown of sheath-promoting proteins attenuates mechanical nociceptive behavior (Jiang et al., 2019). While many studies have focused on how disruption of the integrin pathway alters nociceptor-ECM interactions, the potential role of collagen (a major protein component of ECM) and its receptors in dendrite growth and function remains to be explored. Here, we investigate the role of *smoke alarm* in nociceptor growth and nociceptive behavior. We find that *smoke alarm* promotes dendrite morphogenesis and simultaneously inhibits epidermal ensheathment, and that these functions act to negatively regulate thermal nociception behavior.

## Results

### *smoke alarm* is required for thermal nociception behavior

A tissue-specific RNAi line targeting knockdown of *smoke alarm* in larval nociceptors leads to hypersensitive nocifensive responses to noxious heat and increased dendrite coverage of the larval body wall (Honjo et al., 2016). In order to further investigate the function of *smoke alarm* in nociception, we generated a DNA null mutant allele which removed the entire *smal* gene (Figure 1A) via a FLP-FRT mediated recombination (Parks et al., 2004) between two FRT-containing transposable elements, P{XP}^d03370^ and PBac{WH}^f04782^. Following molecular verification of the breakpoints of the resulting *smoke alarm* deficiency (*smal*^*Df*^) stock, we tested this mutant in our 42°C thermal nociception behavioral assay for hyperalgesia. Consistent with tissue-specific RNAi phenotypes, homozygous *smal*^*Df*^ mutant animals showed hypersensitive thermal nociception responses compared to heterozygous and the genetic background control larvae (Figure S1). The *smal*^*Df*^ animals displayed allodynia relative to the genetic background strain as rolling was observed even with a stimulus temperature that was below the behavioral threshold for nociception responses in this genetic background (42°C). To test for hyperalgesia, we assayed with higher temperature stimuli (46°C probe) in which the background strain control did show nociception behaviors (Figure 1B). *smal*^*Df*^ mutant larvae also showed marked hypersensitivity at 46°C noxious thermal temperature. Interestingly, animals heterozygous for the *smal*^*Df*^ mutation exhibit an intermediate, hypersensitive response to the 46°C stimulus (Figure 1B) further suggesting that nociception sensitivity is sensitive to *smal* gene dosage. Combined, these data indicate both thermal allodynia and hyperalgesia defects for *smal*^*Df*^ null mutant animals.

**Figure 1.**
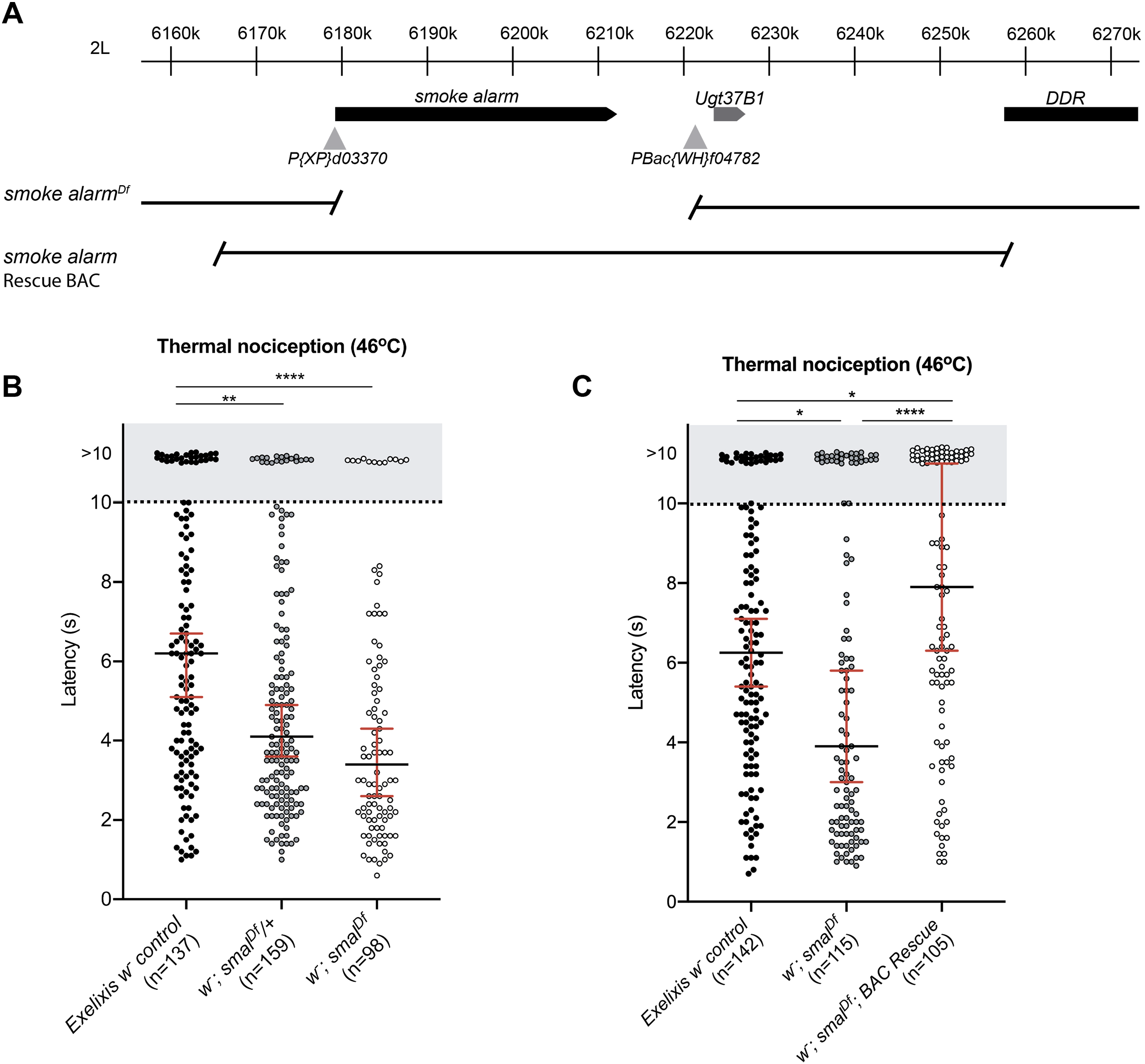
*smoke alarm* mutant larvae show hypersensitivity to noxious heat. **A)** Schematic representation of the *smoke alarm* locus and the *smoke alarm* alleles generated in this study. The *smoke alarm* gene is located on chromosome 2L. The location of the pBac- and P-transposable elements containing FRT sequences used to generate the *smal* null mutant are shown as grey triangles, with the resulting breakpoints of the *smal*^*Df*^ allele denoted. The rescue BAC (Dp(2;3)GV-CH321-55J08) covers the entire genetic interval for *smoke alarm* and its neighboring gene, *Ugt37B1*. **B)** *smal*^*Df*^ null mutant larvae are hypersensitive to noxious thermal stimuli. The rolling latencies of individual larvae presented with a 46°C thermal probe are plotted as distinct points on the graph. Exelixis control larvae (black circles, isogenic *w*^*1118*^, n=137) and larvae that are heterozygous (grey circles, *w*^*1118*^; *smal*^*Df*^/+, n=159) or homozygous (empty circles, *w*^*1118*^; *smal*^*Df*^, n=98) for the *smal*^*Df*^ mutation. Three replicate trials were tested for each genotype. **C)** Genetic rescue experiments of a BAC containing the *smoke alarm* gene in a *smal*^*Df*^ mutant background restore normal thermal nociception responses. The rolling latencies of individual larvae presented with a 46°C thermal probe are plotted as distinct points on the graph. Exelixis control larvae (black circles, isogenic *w*^*1118*^, n=142), homozygous *smal*^*Df*^ mutant larvae (grey circles, *w*^*1118*^; *smal*^*Df*^, n=115), and BAC rescue larvae (empty circles, *w*^*1118*^; *smal*^*Df*^; *DpCH321-55J08V*^*K0003*^*1*, n=105). Three replicate trials were tested for each genotype. Kruskal-Wallis test with Dunn’s test for post-hoc multiple comparisons were performed on data in B and C (*p < 0.05, ***p < 0.001, ****p < 0.0001), only statistically significant comparisons are labeled. Data are presented with median (black bars) ± 95% confidence intervals (red error bars).

To further test whether loss of *smal* (and not an unlinked mutation in the strain) led to the nociception defect, we performed genomic rescue experiments with a chromosomal duplication of wildtype DNA spanning the entire *smoke alarm* locus and its neighboring gene *Ugt37B1* (contained in a 98.2kb bacterial artificial chromosome (BAC), Dp(2;3)GV-CH321-55J08) (Figure 1A). The BAC transgene was crossed into the *smal*^*Df*^ genetic mutant background and tested for phenotypic rescue in our 46°C thermal nociception behavioral assay (Figure 1C). Restoring two copies of *smal* with the BAC fully rescued the hypersensitivity to noxious heat observed in *smal*^*Df*^ mutant larvae (Figure 1C). These rescue results were comparable with behavioral responses to a 46°C stimulus from control larvae. Combined, these data are consistent with the hypothesis that hypersensitive nociception defects in *smal*^*Df*^ mutant larvae are in fact due to the removal of *smal*.

### *smoke alarm* shows expression in class IV md-da sensory neurons

Microarray data from Honjo *et al*. detected an enrichment of *smal* transcripts in larval clV md-da neurons relative to class I md-da neurons (Honjo et al., 2016). However the complete gene expression pattern of *smoke alarm* is not known. Therefore, we used the Trojan-GAL4 system (Diao et al., 2015) to generate an allele of *smal* that expresses GAL4 (Figure 2A, Figure S2A-C). In this allele, an intronic T2A-GAL4 cassette is spliced into the endogenous transcript under control of endogenous regulatory elements and prior studies have shown that this method produces highly accurate reporter gene expression. Consistent with the microarray analyses that showed enriched expression of *smal* RNA in the cIV md-da neurons, this reporter showed robust expression in cIV nociceptive neurons (Figure 2B). We also tested an independently derived *smal*^*GAL4*^ allele that was generated by the *Drosophila* Gene Disruption Project (*smal*^*MI06226-TG4*.*1*^, data not shown) and found a similar pattern (data not shown). In addition, expression was observed in other classes of multidendritic sensory neurons, a subset of type I ciliated neurons, accessory cells (ie, chordotonal cap cells) (Figure 2B) and in motor neurons innervating muscles. Thus *smoke alarm* is indeed expressed in cIV nociceptive neurons of *Drosophila* larvae but it is also expressed elsewhere indicating a potentially more widespread role in the nervous system.

**Figure 2.**
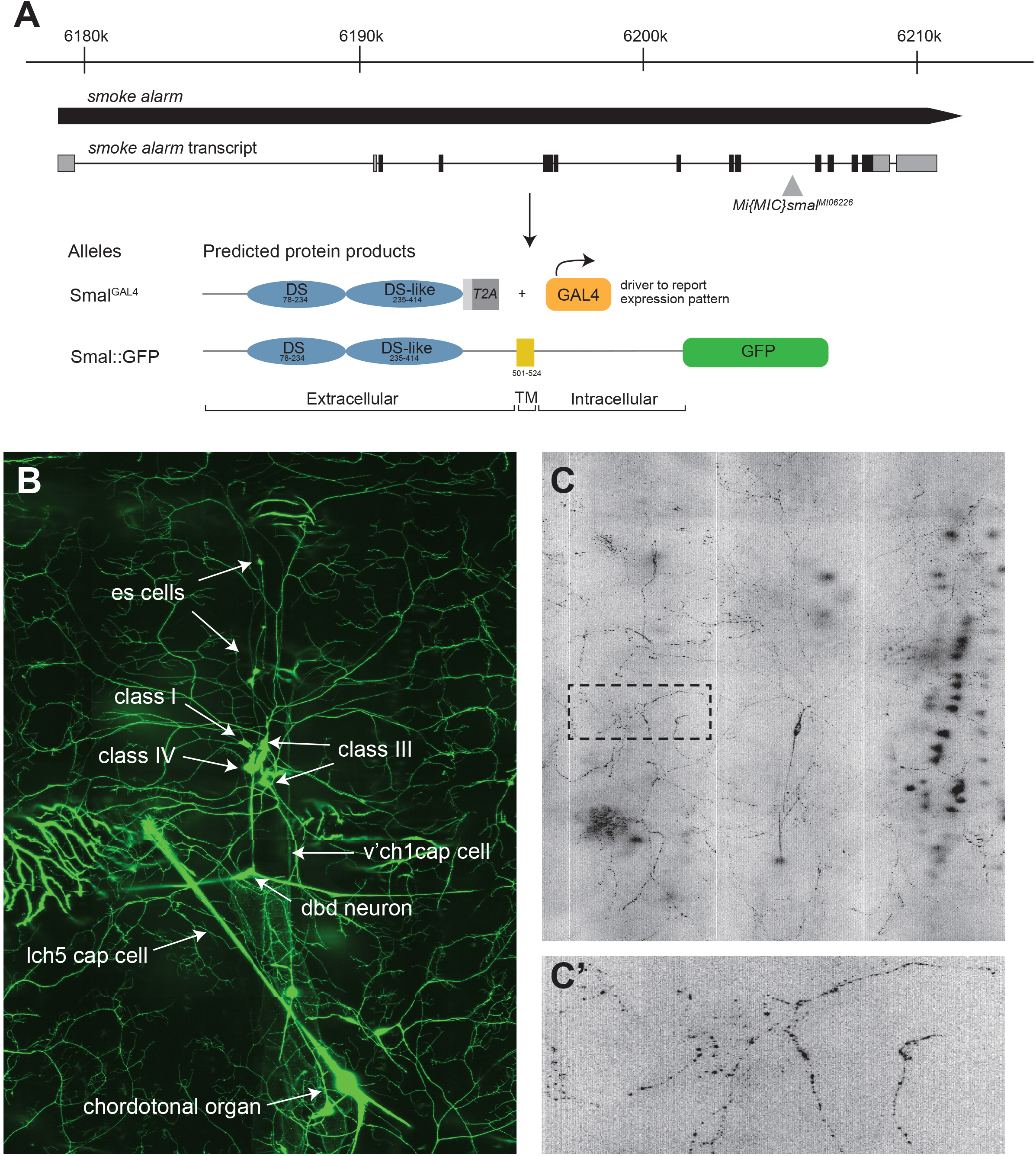
The *smoke alarm* gene expresses in somatosensory md-da neurons. **A)** Schematic representation of the *smoke alarm* gene, transcript, and the expression alleles generated in this study. The *smoke alarm* transcript consists of 10 coding exons that encode a protein with an extracellular discoidin domain (DS, blue oval), a single transmembrane domain (TM, yellow box), and an intracellular region. The *Minos* transposable element (*Mi{MIC}smal*^*MI06226*^, *grey triangle)* located within a coding intron of the *smal* transcript was used to generate the *smal*^*GAL4*^ allele. Predicted protein products from *smal*^*GAL4*^ and *UAS-smal::GFP* alleles are shown indicating location of GAL4 and GFP proteins. Numbers within the domains denote the relevant amino acid residues within the predicted protein products. **B)** *smoke alarm* is expressed in neurons of the larval peripheral nervous system. A confocal micrograph of the *smal*^*Gal4*^ gene expression pattern centered on the dorsal cluster and part of the lateral cluster of segment A5 in a wandering third instar larva. Left is posterior, dorsal is up (*w*^*1118*^; *smal*^*GAL4*^*/UAS-mCD8::GFP*^*LL4*^). GFP-postiive signals were detected in each larval segment along the body wall. *smal*^*Gal4*^ *i*s expressed in a subset of type I ciliated neurons (external sensory neurons (desA-D) and chordotonal organs (v’ch1 and lch5)) and type II non-ciliated multidendritic neurons (class I (ddaD and ddaE), class III (ddaA and ddaF), class IV (ddaC), and the dorsal bipolar dendrite (dbd) neurons) in the peripheral nervous system. Additional GFP labeling of accessory cap cells. **C)** Smal::GFP proteins localize along dendritic arbors of class IV neurons. A confocal micrograph of ddaC (*w*^*1118*^; *ppk-GAL4/ UAS-smal::GFP*) shows detectable levels of GFP in the soma, axon, and along the dendritic arbor.

We additionally explored *smal*^*GAL4*^ driven mCD8::GFP expression in the larval brain and observed expression in regions of the central brain, the developing optic lobe, and the ventral nerve cord (VNC; Figure S2D). This expression included neurons in different compartments of the VNC and glial cell populations throughout the brain. In order to tease apart *smal*^*GAL4*^ expression in potential subtypes of larval brain tissue, we examined animals with *nsyb-GAL80* and *tsh-GAL80* repressors in combination with the *smal-GAL4>UAS-mCD8::GFP* reporter. The inhibition of *smal*^*GAL4*^ expression with the *nsyb-GAL80* repressor that targets late born neurons in the CNS showed reduced GFP-labeled neuronal cell bodies in the VNC (Figure S2D) while repression experiments with *repo-GAL80* led to a complete loss of GFP-labeled cells on the surface of the brain lobes and the VNC (Figure S2D). Combined these data reveal that the *smoke alarm* gene is expressed in a population of neurons and glial cells in the central nervous system, which is in contrast to neuronal *smal* expression in the peripheral nervous system where glial expression was not observed.

### Smoke Alarm proteins localize to nociceptor dendrites

The Smoke Alarm (Smal) protein is a *Drosophila* homologue of Discoidin Domain Receptors which have extracellular domains that bind collagen in the extracellular matrix and previous studies have found the cIV md-da neuron dendrites adhere to the extracellular matrix at the basolateral surface of epidermal cells. Given this, and due to the dendrite phenotypes of *smal*^*Df*^ larvae, we hypothesized that Smal might localize to dendrites. Consistent with this hypothesis, we found that a c-terminally GFP tagged Smal protein (Smal::GFP) (Figure 2A) expressed under control of the GAL4-UAS system was localized to the cIV neuron dendrite arbor of third instar larvae (Figure 2C). At higher magnification, GFP-tagged Smal proteins show a punctate distribution pattern throughout the dendritic arbor to the distal ends (Figure 2C’). Given the known function of mammalian DDR proteins, these sites of Smal::GFP may be indicative of regions where dendrites attach to collagen in the basal lamina. Importantly, the presence of Smal::GFP in dendrites is consistent with the hypothesis that Smal functions in regulating dendrite morphology of cIV neurons. However, the Smal::GFP proteins were also detected in the cell body and in the axon, including its terminals which raises the possibility the Smal protein functions in more than one neuronal compartment.

### *smoke alarm* mutants display defects in nociceptor dendrite morphogenesis

A *UAS-smal-RNAi* line under control of a cIV da-GAL4 esulted in a dendrite hyperbranching phenotype (Honjo et al. 2016). This suggested the possibility that *smoke alarm* may regulate dendrite morphogenesis of cIV neurons. To investigate this hypothesis further, we imaged the dendritic field of cIV neurons in *smal*^*Df*^ mutant larvae and traced their arbors (Figure 3A, Figure S3A-C). In contrast to the RNAi phenotype for *smal*, null mutation for *smal* resulted in a significant decrease in dendrite arbor coverage. Interestingly, larvae deficient for *smoke alarm* showed a reduction in total dendritic length when compared to control animals while heterozygous animals showed an intermediate reduction in total dendritic length (Figure S3D). This difference in dendritic length persisted when corrected for neuron area (Figure 3B). Additionally, *smal*^*Df*^ mutants exhibited a mild reduction in the dendritic area and the number of dendrite terminals of cIV neurons (Figure S3D). Combined, nociceptive neurons in *smal*^*Df*^ larvae show a less complex dendritic arborization pattern, suggesting that the wild type *smoke alarm* gene function promotes dendritic growth and branching of cIV neurons. This whole animal mutant phenotype contrasts with previously described RNAi targeting *smal* specifically in cIV neurons which caused hyperbranching.

**Figure 3.**
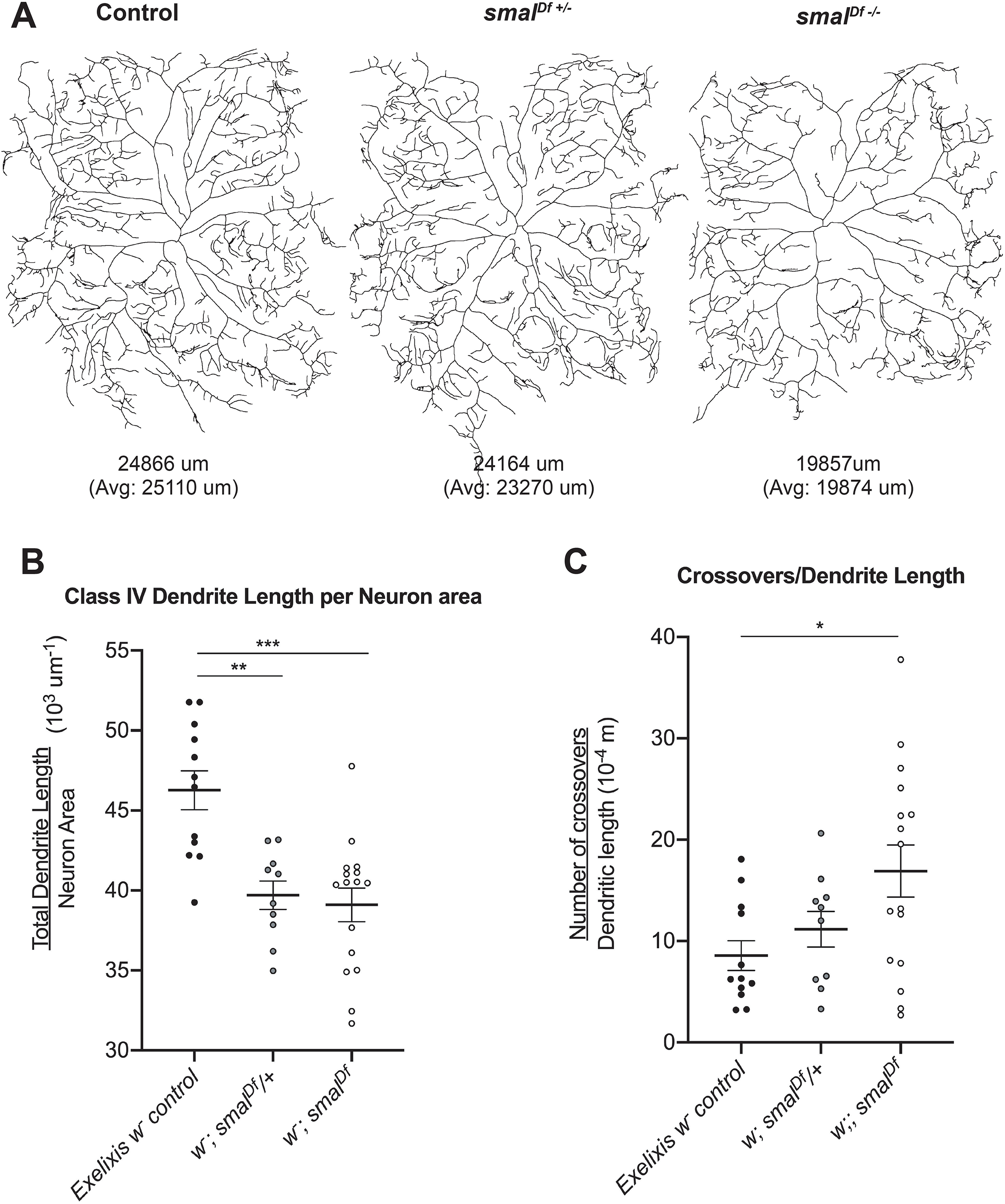
*smoke alarm* mutant larvae display defects in nociceptor dendrite morphogenesis. **A)** Class IV neurons in *smoke alarm* mutant larvae show reduced dendritic length. 2-D neuronal reconstructions of the dorsal class IV neuron, ddaC, from confocal micrographs from third instar wandering larvae. Left is posterior, dorsal is up. Representative tracings are shown for control larvae (*w;; ppk-CD4::tdTomato/+*), and animals heterozygous (*w; smal*^*Df*^*/+; ppk-CD4::tdTomato/+*) or homozygous (*w; smal*^*Df*^; *ppk-CD4::tdTomato/+*) for the *smal* deficiency mutation. The corresponding total dendrite length (um) is indicated below each genotype’s representative tracing. **B)** Class IV neurons in *smoke alarm* mutant larvae show a decrease in total dendrite length per neuronal surface area. Analysis and quantification of total dendrite length corrected for total neuron area in Exelixis w^-^ control (n= 12), *smal*^*Df*^*/+* (n=10), and *smal*^*Df*^ (n=16) mutant class IV neuron tracings. **C)** Class IV neuron dendrites in *smoke alarm* mutant larvae show an increase in isoneural crossing events. Analysis and quantification of the number of crossovers standardized to total dendritic length in Exelixis *w*^*-*^ control (n= 12), *w*^*-*^; *smal*^*Df*^*/+* (n=10), and *w*^*-*^; *smal*^*Df*^ (n=16) mutant class IV neuron tracings. Welch’s ANOVA test with Dunnett’s test for post-hoc multiple comparisons (*p < 0.05, **p < 0.01, ***p < 0.001) were performed on data in B and C. Data are presented as mean (black bar) ± SEM (error bars).

### *smoke alarm* mutants show an enhanced epidermal ensheathment phenotype

Several studies have reported that adhesion of nociceptive dendrites to the ECM of the basal lamina via integrin-laminin interactions is critical for dendritic morphogenesis, patterning, and function (Han et al., 2012; Kim et al., 2012). Given that mammalian DDR proteins bind to collagen we hypothesized that Smal may be similarly important in adhesion of dendrites to the basal lamina and that loss-of-function *smal* mutants might show aberrant ECM attachment. Consistent with this hypothesis, we observed an increased in branch crossovers for isoneural dendrites of cIV neurons from *smal*^*Df*^ mutant larvae (Figure 3C and Figure S3D). We also observed increased crossing events in animals with cIV md-da specific RNAi knockdown of *smal*. A similar dendrite crossing phenotype has been described in studies of integrin loss-of-function in cIV md-da neurons where it has been shown that crossing over of dendrites occurs when they detach from the ECM and become enveloped by the epidermis (Han et al., 2012; Kim et al., 2012). Close inspection of confocal z-stacks indicate that the crossing events we observed in cIV neurons of *smal*^*Df*^ mutant larvae were in non-contacting dendrites in distinct focal planes (data not shown).

Thus, we investigated whether *smal* mutant dendrites showed an increase in ensheathment by overlying epidermal cells. To test this, we investigated a marker for epidermal ensheathment (the pleckstrin homology domain reporter, PLCδ-PH-GFP). This marker shows localization to the epidermal plasma membrane in regions enriched for phosphatidylinositol 4,5-bisphosphate (PIP2) which in turn correlates with areas of dendritic ensheathment (Jiang et al., 2019) (Figure 4A, 4B, 4B’). By co-labeling the plasma membrane of cIV neuron dendrites with a tandem dimer Tomato florescent protein (tdTomato; Figure 4A, 4B, 4B’’), we traced the entire cIV dendritic arbor and measured the portion that co-localized with PIP2-marked ensheathment (Figure 4A, 4B, 4B’’’, 4C). Larvae deficient for *smoke alarm* showed a significant increase in total ensheathed dendritic length when compared to control or heterozygous animals (Figure 4D). Although percent ensheathment increased in the *smal*^*Df*^ mutant, Sholl analysis did not uncover any changes in the spatial domain of the epidermally-enclosed dendrites. In summary, we observe both an increase in dendrite crossing events as well as increases in ensheathment in *smal*^*Df*^ animals. Thus, we propose that Smoke Alarm proteins function as dendrite ECM adhesion molecules and that loss of such adhesion leads to hyperalgesia.

**Figure 4.**
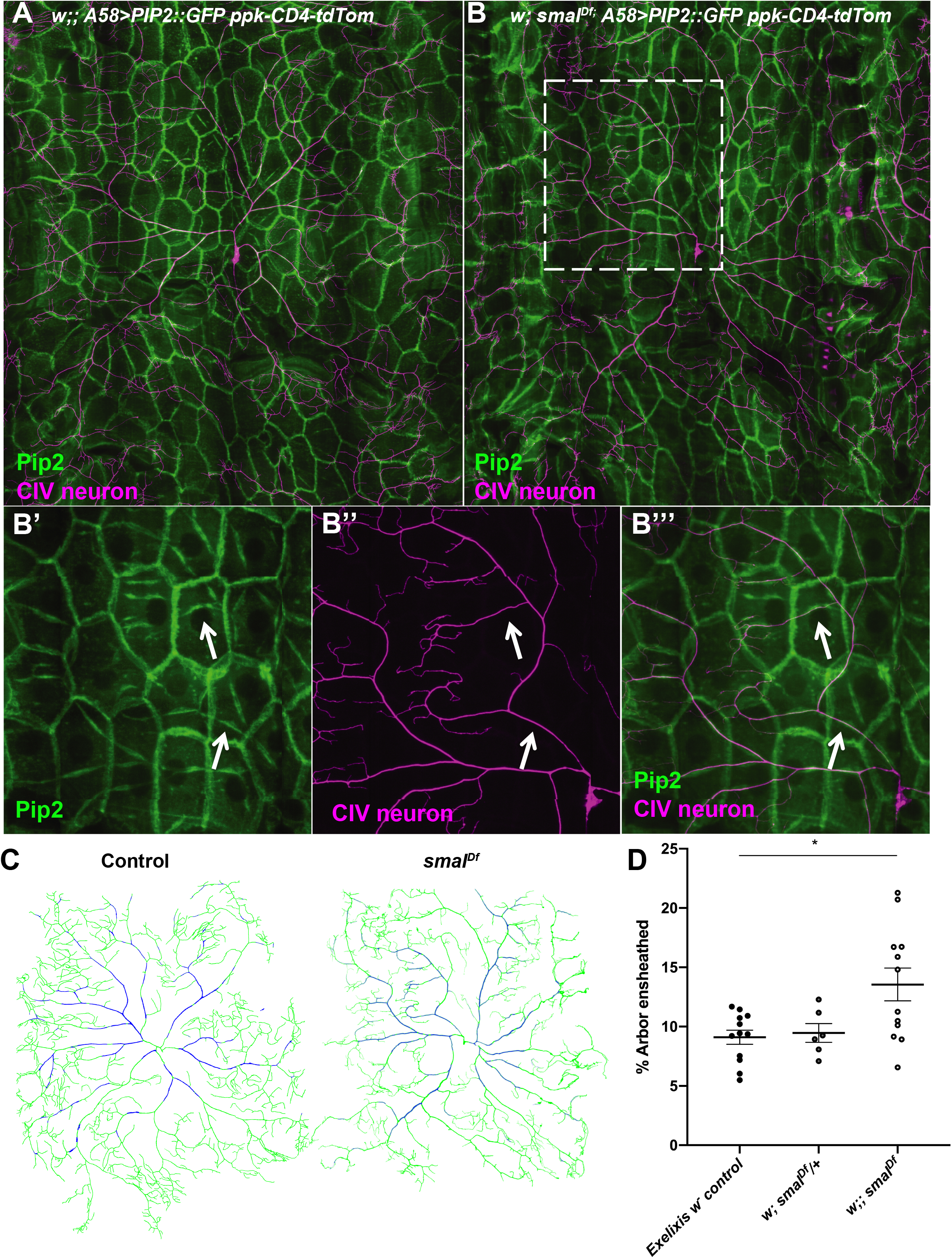
*smoke alarm* mutants show mild epidermal ensheathment phenotypes of Class IV dendrites. **A-B)** Class IV neurons in *smoke alarm* mutant larvae show increased epidermal enclosures of nociceptor dendrites. Epidermal expression of GFP-tagged PIP2 labels ensheathment of class IV dendrites into the epidermis. *A58-GAL4* expresses PLCδ-PH::GFP (green) in the epidermis of larvae co-expressing a membrane-targeted marker in class IV-neurons (magenta, *ppk-CD4-td::Tomato*) of control larvae. Representative confocal micrographs of class IV ddaC neurons from wandering third instar larvae are shown for A) control and B) *smal*^*Df*^ mutant. High magnification images (B’, B’’, and B’’’) of inset (B) show PIP2 accumulation (green, B’ and B’’’) at ensheathed class IV dendrites (magenta, B’’ and B’’’). **C)** 2-D neuronal reconstructions from confocal micrographs of the dorsal class IV neuron, ddaC, third instar wandering larvae. Left is posterior, dorsal is up. Representative tracings are shown. Green represents all of the traced dendrites and blue denotes the portion of dendritic arbor that is ensheathed by the overlying epidermis. **D)** Class IV neurons in *smoke alarm* mutant larvae show an increase in the percent of dendrite arbor that is ensheathed. Analysis and quantification of the proportion of ensheathed dendrite relative to the entire dendrite length in class IV neuron tracings from control larvae, and animals heterozygous or homozygous (for the *smal* deficiency mutation. Welch’s ANOVA test with Dunnett’s test for post-hoc multiple comparisons was performed. Data are presented as mean ± SEM, *p < 0.05.The genotypes and number of neurons used in the quantification and analysis of dendrite morphogenesis parameters were *w;; A58-GAL4 UAS-PLCd-PH::EGFP/ppk-CD4::tdTomato* (control larvae, n=12 neurons), *w; smal*^*Df*^*/+; A58-GAL4 UAS-PLCδ-PH::EGFP/ppk-CD4::tdTomato* (heterozygous mutant larvae, n=6 neurons), and *w; smal*^*Df*^; *A58-GAL4 UAS-PLCd-PH::EGFP/ppk-CD4::tdTomato* (homozygous mutant larvae, n=12 neurons).

## Discussion

Taken together, our data suggest that *smoke alarm* broadly regulates class IV da neuron morphogenesis and function. With *smal* loss of function, several distinct and potentially related cellular phenotypes arise in cIVda neurons. First, dendritic arbors are smaller and they show less branching relative to animals with wild type *smal* function. Second, *smal* mutant isoneural dendrites show an increased tendency to cross over one another. Third, *smal* mutant dendrites show an increase in ensheathment by overlying epidermal cells. Finally, in behavioral assays, *smal* mutant larvae show both thermal allodynia and hyperalgesia phenotypes.

These data combined raise interesting questions for future study. For instance, are the morphological phenotypes a consequence of neuronal hypersensitivity? Or alternatively, does Smal function regulate dendrite morphology which in turn results in hypersensitivity? We favor the latter hypothesis. Smal::GFP proteins localize to the dendritic arbor and it contains a Discoidin Domain which is predicted to bind collagen. Thus, Smal protein is in the correct place to be involved in dendrite morphogenesis and it possesses a known adhesion domain. This adhesion function may be needed for anchoring dendrites to the ECM as they grow, or it may be needed for the stabilization of dendritic branches that have already grown.

Whether or not the dendrite phenotypes that we observe contribute to nociceptor hypersensitivity also remains unknown. In our prior studies, a phenotype of reduced branching in nociceptor dendrites is correlated with insensitive phenotypes (Honjo et al., 2016). In the case of *smal*, hypersensitivity phenotypes are observed even though dendritic arbors and branching are reduced. Perhaps this is explained by the fact that *smal* mutants show a relative increase in epidermal ensheathment even with correction for reduced arbor size. Thus, an increased interaction with epidermal cells may lead to hypersensitivity that is epistatic to an insensitivity that is caused by reduced branching. Consistent with this hypothesis, mammalian keratinocytes have been shown to signal to nociceptive nerve endings and to facilitate the excitability of nociceptors (Baumbauer et al., 2015; Pang et al., 2015). The mutant phenotype of *smal* suggests that similar epidermal signals may exist in *Drosophila*. Indeed, previous studies have found that larval epidermal cells express the TGF-beta ligand Maverick which signals to the cIVda neurons through the receptor tyrosine kinase Ret to promote dendrite growth (Hoyer et al., 2018). Interestingly, Ret has also been proposed to regulate dendrite adhesion through an interaction with integrins and Rac1 signaling (Soba et al., 2015).

Although the focus of our study has been on the nociceptor dendrites and epidermal interactions, it is interesting to note that Smal::GFP is also detected in axons including the axonal regions that contain nociceptor synapses. Prior studies in *Drosophila* have found a variety of important function for ECM and proteins that interact with ECM at synapses (Rohrbough et al., 2000; Rushton et al., 2009). Thus, although we have not detected any gross morphological defects in cIV axonal projections in *smal* mutant background, it remains a possibility that Smal function at synapses could affect excitability of nociceptors and other neurons that express the gene.

Smal is homologous to Mammalian Discoidin Domain Receptor proteins, which possess a Receptor Tyrosine Kinase activity that is activated by a collagen ligand. The intracellular domain (ICD) of Smal protein itself is unlikely to possess RTK activity as it lacks the highly conserved HRD and DFG motifs that are diagnostic of kinase domains. Thus, Smal may function primarily in adhesion, but it remains possible that the Smal intracellular domain has signaling activity. Future structure-function studies (ie. functional rescue tests with constructs lacking the ICD) are needed to address this possibility.

## Acknowledgements

Stocks obtained from the Bloomington Drosophila Stock Center (NIH P40OD018537) and the Exelixis Collection Harvard Medical School were used in this study. We thank Tzumin Lee (Janelia Research Campus) for constructing the *repo*-GAL80 transgene and for sharing it with us. We thank James Truman (Janelia Research Campus) for generating and sharing the *nsyb*-GAL80 transgene. We also thank Luuli Tran, Lauren Eads, and Rebecca McAvoy for technical assistance and members of the Tracey lab for helpful discussion. Research reported in this publication was supported by the National Institute of General Medical Sciences of the National Institutes of Health under award number R01GM086458.

## Experimental Procedures

### Fly Strains and Husbandry

All larvae used in experiments were reared on the Bloomington *Drosophila* cornmeal medium in a 25**°**C incubator with controlled humidity (70%) on a 12h light/12h dark cycle. Strains were otherwise maintained at room temperature. The following strains were used in this study: *Exelixis w*^*1118*^ ((Parks et al., 2004) BDSC #6326), *Df(2L)ED385* ((Ryder et al., 2004)BDSC 9341), *yw; Mi{MIC}smal*^MI06226^ ((Venken et al., 2011) BDSC #40801), *w;; ppk-CD4::tdTomato* (Han et al. (2011)BDSC #35845), *yw;{UAS-mCD8::GFP}LL5* ((Lee & Luo, 2001) BDSC #5139), *Mi{Trojan-GAL4*.*1}smal*^*MI06226- TG4*.*1*^ (BDSC #67464), *yw;; UAS-PLCδ-PH-eGFP* (PIP2, BDSC #39694), *GMR57C10-GAL80 (nsyb-GAL80*) (Harris et al., 2015), *w; repo-GAL80 (Awasaki et al*., *2008), w; ppk1*.*9-GAL4* (Ainsley et al., 2003), *A58-GAL4* (Galko & Krasnow, 2004).

#### Generation of smal^Df^ allele (this study) and BAC rescue strain

Exelixis lines, *P{XP}d03370* and *PBac{WH}f04782*, were provided by the Exelixis collection at Harvard Medical School (Thibault et al., 2004) and used to generate FLP-FRT mediated *smal*^*Df*^ mutant flies (crossing strategy outlined in (Parks et al., 2004)). Candidates were crossed to the *w; ED385* deficiency and molecularly verified by PCR using primers that amplify the hybrid element following recombination (‘5-AATGATTCGCAGTGGAAGGCT-3’ and 5’-GACGCATGATTATCTTTTACGTGAC-3’) and primers that target the *smoke alarm* locus (primer pair 5’-GCAGCGTTTCGTTTGCCG-3’ and 5’-GTCGCCAGTACTCCAGGACATAG-3’, primer pair 5’-TGTGGACGACAACGATTCG-3’ and 5’-TGCGGAAGTTGTAGGTCCTG-3’). The *smal* genomic rescue strain utilized the interchromosomal duplication line *w;;Dp(2;3)GV-CH321-55J08*^*VK00031*^ from GenetiVision Corporation. Transgenic flies containing the BAC rescue construct were crossed into the *smal*^*Df*^ deletion background.

#### Generation of Trojan-Gal4 allele (this study)

We generated the *smal*^*GAL4*^ allele according to methods described in Diao *et al*., (2015). The triple phase Trojan-GAL4 donor cassette, *yw; Sp/CyO; P{lox(Trojan-GAL4)x3}11*) ((Diao et al., 2015) BDSC #60311), and the *smal* insertion line containing the intronic *Mi{MIC}*^*MI06226*^ element ((Venken et al., 2011) BDSC #40801) were used for recombinase-mediated cassette exchange to generate *smal*^*GAL4*^ flies (crossing strategy outlined in (Diao et al., 2015)). Orientation of the inserted cassette and the correct linker (phase 1) was confirmed with PCR amplification and sequencing (Figure S2). Primers used to amplify the upstream region (primer pair 1 in Figure S2) are 5’-GATTTGCTCGCGGTGATTG-3’ and 5’-CGCTATCGATGCTCACGGTC-3’ (expected product size 1320bp). The primers used to amplify the downstream region (primer pair 2 in Figure S2) are 5’-CCAACTTCAACCAGAGCGG -3’ and 5’-GTTTCGACACTGCCATTG-3’ (expected product size 1590 bp).

#### Generation of UAS-smal::GFP (This study)

Model System Injections performed microinjections of *pUAST-smal::GFP* vectors into *w*^*1118*^ and transgenic flies were generated via standard P-element mediated transformation.

### Behavioral Assay

Thermal nociception behavior experiments were performed as previously described in (Caldwell & Tracey, 2010; Honjo et al., 2016). Larvae were assayed for hypersensitivity at 42**°**C (variac set at 21V) to test for allodynia and 46**°**C (variac set at 23V) to test for hyperalgesia. Crosses were set up with 6 females and 3 males per vial on standard cornmeal medium (BDSC recipe). Crosses were kept in an incubator set to 25**°**C at 70% humidity. Parental flies were removed 5 days following mating and larvae were tested 6-7 days after the initial cross was set up. Experiments were performed in triplicate.

### Immunohistochemistry

Larval brains were dissected and immunostained according to standard protocols (detailed protocols are available upon request). The following antibody reagents were used for immunofluorescence at the following concentrations: rabbit anti-GFP (1:500, Thermo Fisher Scientific); mouse anti-nc82 (1:30, Developmental Studies Hybridoma Bank); Alexa Fluor 488 goat anti-rabbit (1:1000, Thermo Fisher Scientific); Alexa Fluor 635 goat anti-mouse (1:1000, Thermo Fisher Scientific).

### *In vivo* Confocal Microscopy

To observe dendrite morphology of md-da neurons in live imaging experiments, we lightly anesthetized wandering 3^rd^ instar larvae with diethyl ether for 15 minutes in a glass chamber. Larvae were then mounted in glycerol on a glass microscope slide under a 22×50mm #1.5 coverslip. For consistency, only sensory neurons from the dorsal cluster located in segments A4-A6 were imaged using the tile scan feature on a confocal LSM Zeiss 5 Live with a 40x apochromat immersion oil objective. Confocal Z-stacks were converted to maximum intensity projections using the Zeiss Zen software package. Negative image conversion and adjustments to brightness and contrast were performed in Adobe Photoshop.

### Dendrite Morphology Analysis

Confocal micrograph Z-stacks of class IV ddaC dendritic arbors were traced as maximum intensity projections using the Simple Neurite Tracer plug-in (Longair et al., 2011) for FIJI (Schindelin et al., 2012). Neuron area measurements and the quantification of total dendrite length and number of dendritic terminals were calculated using the Simple Neurite Tracer plug-in. Crossovers were manually counted for each neuron. For ensheathment experiments, only the portion of the dendrite that co-localized with the epidermal expressed PIP2 GFP marker were traced.

### Molecular Biology

*smoke alarm* cDNA was synthesized by GeneArt Gene Synthesis (Thermo Fisher) and cloned into the pTW (*pUAST-smal::GFP*) destination vector (Murphy, 2003) via Gateway recombination (Petersen & Stowers, 2011).

### Experimental design and statistical analysis

For all experiments, controls and genetically matched experimental genotypes were tested at the same time. Mixed sexes were randomly tested in all larval experiments and sample genotypes were blinded during both data acquisition and analysis. The number of animals tested or neurons analyzed is indicated in the figure legends. All statistics were performed using GraphPad Prism 8. Datasets were tested for normality using Shapiro-Wilks goodness of fit tests. Details on statistical tests are provided in figure legends.

