## Supplemental Figures for "The *Drosophila* gene *smoke alarm* regulates nociceptor-epidermis interactions and thermal nociception behavior"

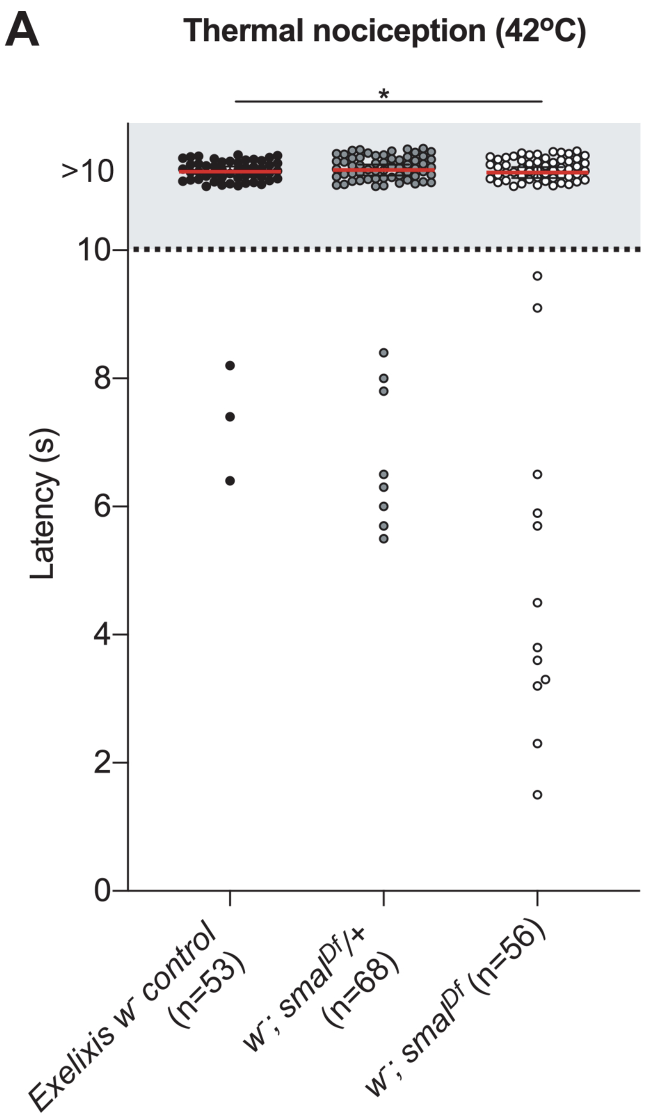


**Supplementary Figure 1**

**Supplemental Figure 1. Smoke alarm mutant larvae show thermal allodynia.**

A) *smal*$^{Df}$ null mutant larvae are hypersensitive to thermal stimuli. The rolling latencies of individual larvae presented with a 42$^{o}$C thermal probe are plotted as distinct points on the graph. Any larvae that did not respond to the stimulus, or took longer than the10s cut-off to roll, were grouped together. Exelixis control larvae (black circles, isogenic *w*$^{1118}$, n= 53) and larvae that are heterozygous (grey circles, *w*$^{1118}$; *smal*$^{Df}$/+, n=68) or homozygous (empty circles, *w*$^{1118}$; *smal*$^{Df}$, n=56) for the *smal*$^{Df}$ mutation. Kruskal-Wallis test with Dunn's multiple comparison tests (*p < 0.05), only significant comparisons are labeled. Red bar denotes the median response latency (>10s). Two replicate trials were tested per genotype.

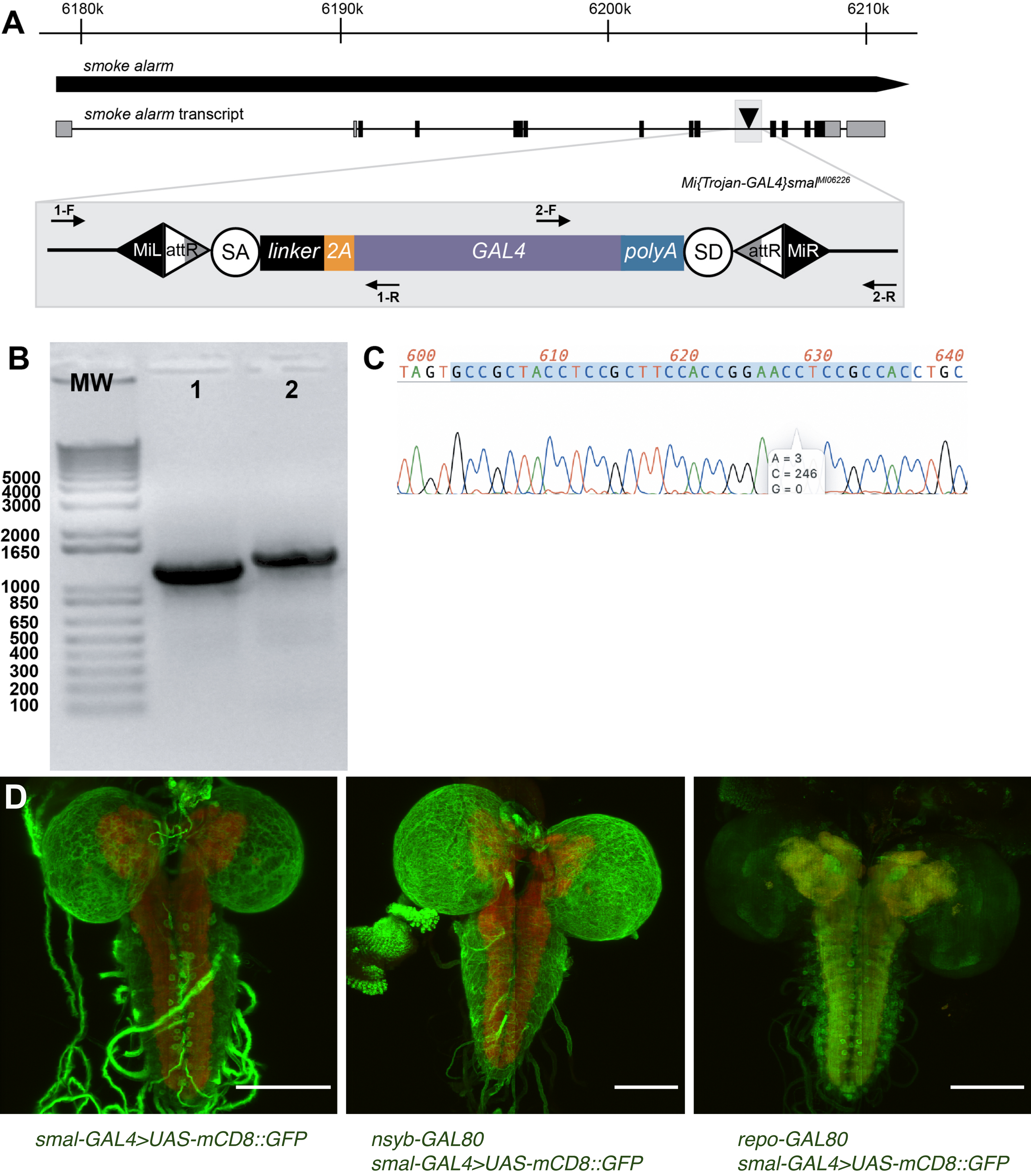


### Supplementary Figure 2

**Supplemental Figure 2. Molecular verification of the *smal*$^{GAL4}$ allele and its CNS expression.**

A) Schematic representation of the *smoke alarm* locus denotes the intronic MiMIC transgene (black triangle) used to generate the *smal*$^{GAL4}$ allele (*Mi{Trojan-GAL4}smal*$^{MI06226}$) in this study. Enlarged view of this region shows the integration of the Trojan-GAL4 exon into the *smal*$^{MI06226}$ element following recombinase-mediated cassette exchange. The resulting *smal*$^{GAL4}$ allele consists of two *Minos* inverted repeats (MiL and MiR) that flank two recombination sites (attR). Within these attR sites lies the Trojan-GAL4 exon cassette that consists of a splice acceptor (SA) site, a short flexible linker, a T2A sequence, the entire GAL4 coding sequence, a Hsp70 polyadenylation (polyA) signal sequence, and a splice donor (SD) sequence. The arrows represent the location of the two primer pairs (1 and 2) used for PCR confirmation experiments (primer set 1: 1120 bp, primer set 2: 1310 bp). B) Agarose gel electrophoresis of PCR amplified products using *smoke alarm* and GAL4 specific primer pairs 1 and 2. The molecular weight (MW) lane indicates the Invitrogen 1kb Plus DNA ladder. The *smal*$^{GAL4}$ animals show amplified products for both primer pairs, denoting Trojan-GAL4 exon insertion with correct orientation. The PCR product from primer pair 1 was sequenced to determine the linker phase. C) Chromatogram of linker phase sequence highlighted in blue confirms a splice phase 1 for the Trojan-GAL4 insertion at the *Mi{MIC}*$^{MI06226}$ site. D) *smal*$^{GAL4}$ expresses mCD8::GFP under the control of *smoke alarm* endogenous regulatory elements and labels neurons and glia of the central nervous system (left). Representative confocal micrographs of dissected brains from wandering third instar larvae are shown. Gene expression is restricted from CNS neurons using *nsyb-GAL80* (middle) or CNS glia using *repo-GAL80* (right). Green represents anti-GFP immunoreactivity against mCD8::GFP. Red represents anti-bruchpilot (nc82) immunoreactivity. Scale bars, 100um. Genotypes were *w*; *smal*$^{GAL4}$ *UAS-mCD8::GFP/*+ (left), *nsyb-GAL80 w/*+; *smal*$^{GAL4}$ *UAS-mCD8::GFP* (middle)*, w*; *smal*$^{GAL4}$ *UAS-mCD8::GFP /repo-GAL80* (right).

**A** Control: w⁻;; *ppkCD4::tdTomato/+*

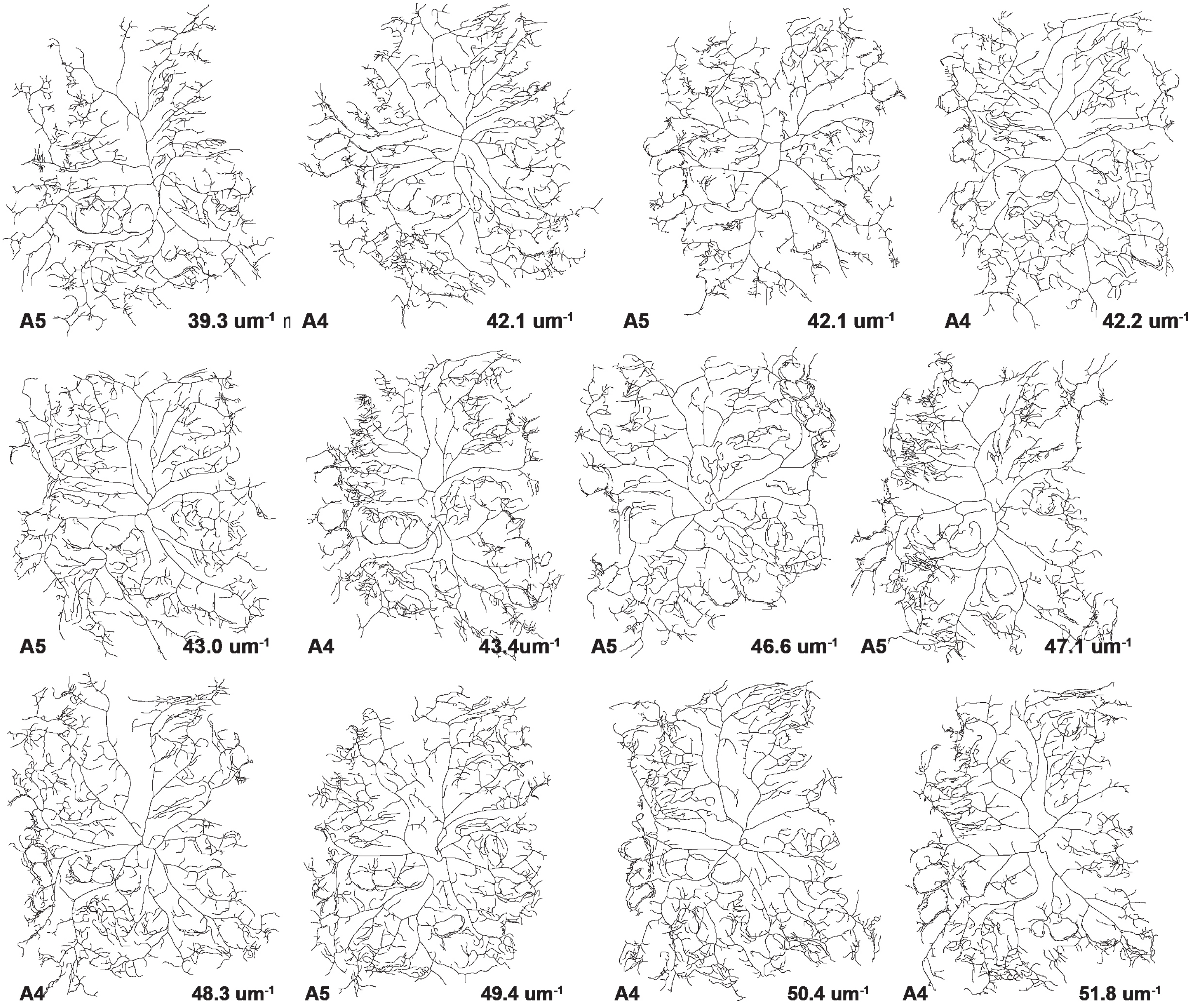


### Supplementary Figure 3A

**B** Heterozygous *smal* mutant larvae: *$w^{-}$; $smal^{Df}$/+; ppkCD4::tdTomato/+*

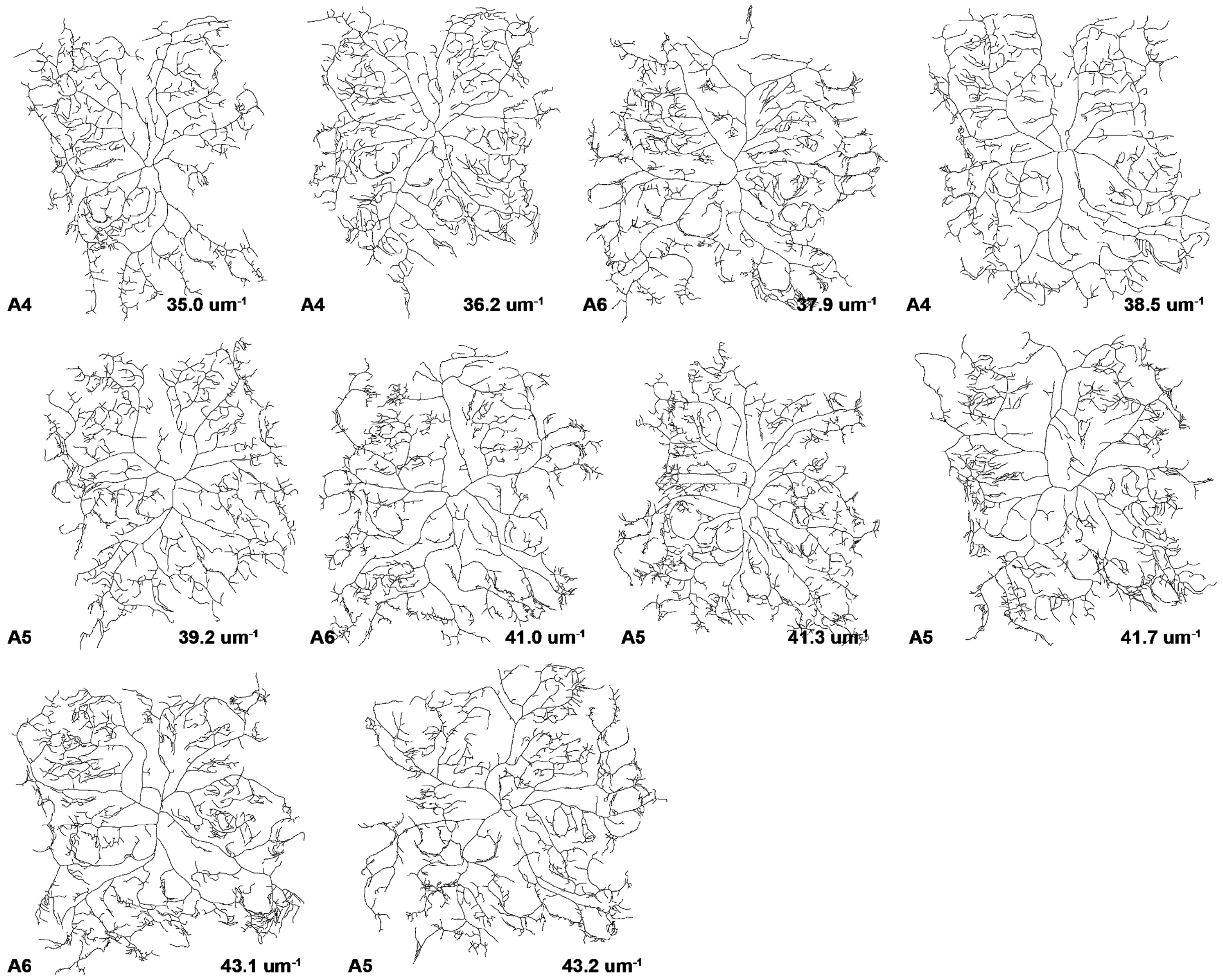


### Supplementary Figure 3B

**C** Homozgous *smal* mutant larvae: $w^-$; *smal*$^{Df}$; *ppkCD4::tdTomato/+*

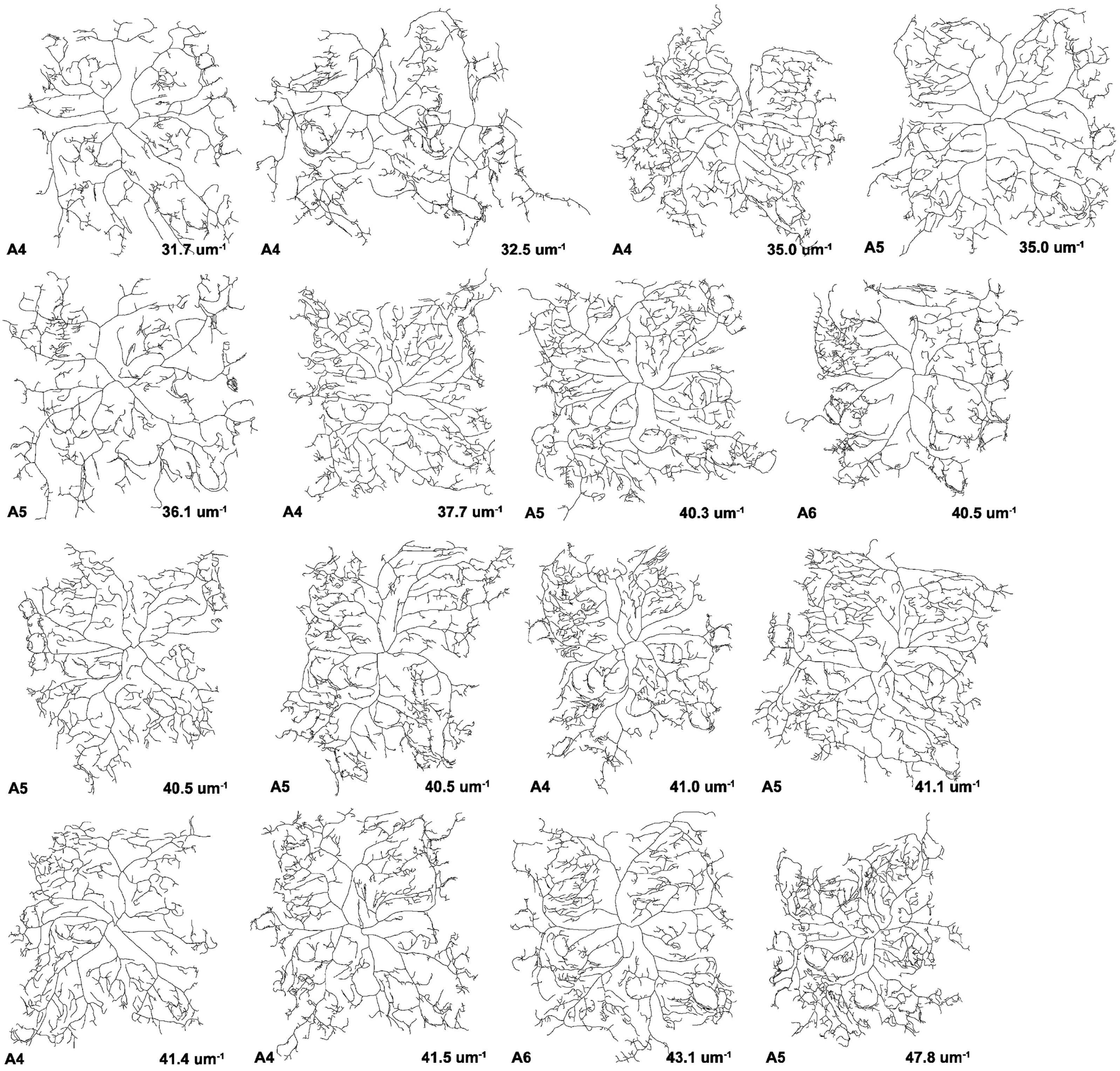


### Supplementary Figure 3C

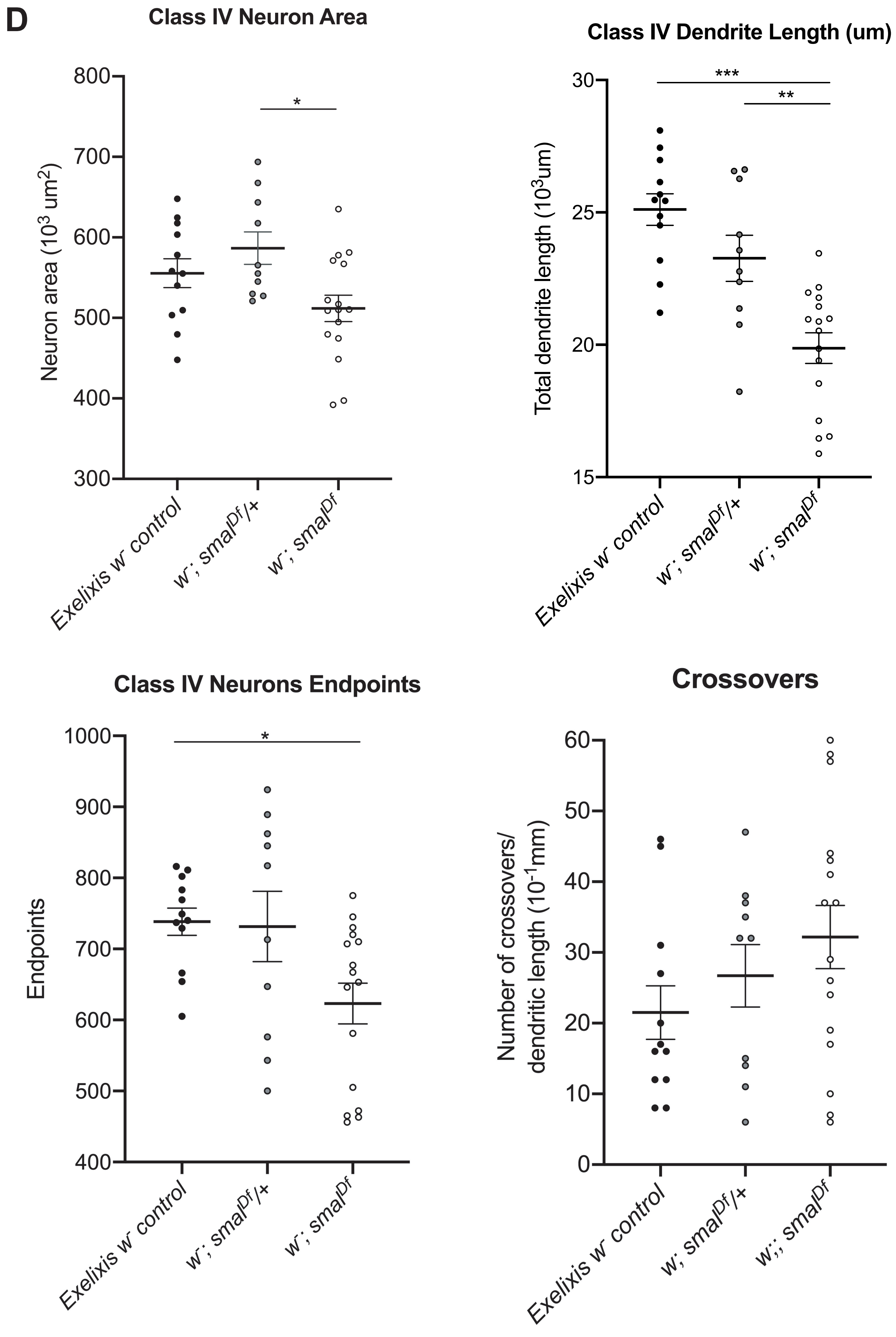


Supplementary Figure 3D

**Supplemental Figure 3. class IV md-da neuronal reconstructions from *smal* mutant larvae.**

2-D neuronal reconstructions of confocal micrographs of the dorsal class IV neuron, ddaC, from third instar wandering larvae. All tracings are shown for A) control larvae (*w;; ppk-CD4::tdTomato/*+), and animals B) heterozygous (*w; smal$^{Df}$/+; ppk-CD4::tdTomato/*+) or C) homozygous (*w; smal$^{Df}$; ppk-CD4::tdTomato/*+) for the *smal* deficiency mutation. The corresponding abdominal segment (A4, A5, or A6) segment for each traced ddaC neuron and its total dendrite length corrected for area ($10^3$ $um^{-1}$) is indicated below each tracing. Left is posterior, dorsal is up. D) Analysis and quantification of total dendrite length, total neuron area, number of endpoints, and number of crossovers from class IV neuron tracings in control (n= 12), *smal$^{Df}$/*+ (n=10) and *smal$^{Df}$* (n=16) mutant class IV neuron tracings. Welch's ANOVA test with Dunnett's test for post-hoc multiple comparisons was performed. Data are presented as mean $\pm$ SEM, *$p < 0.05$, **$p < 0.01$, ***$p < 0.001$.
